# Active Learning for Protein Structure Prediction

**DOI:** 10.1101/2025.04.01.646404

**Authors:** Zexin Xue, Michael Bailey, Abhinav Gupta, Ruijiang Li, Alejandro Corrochano-Navarro, Sizhen Li, Lorenzo Kogler-Anele, Qui Yu, Ziv Bar-Joseph, Sven Jager

## Abstract

Accurate protein structure prediction is challenging especially for families and types that are not well-represented in current training data. Active learning selects candidates for labeling with the aim of most rapidly improving model performance. In a general, many labeling strategies have been proposed. However, with protein structure prediction, most of these strategies don’t apply or are difficult due to the high-dimensional, variable-dimensional regression target and the inherent complexity of the models involved. We applied a novel active learning strategy, DEWDROP, to protein structure prediction on two different protein datasets: VHH-only antibodies (Nanobodies™), and Mycobacterial proteins. We introduce a domain-specific fine-tuned Equifold model for VHH structures and apply DEWDROP to generate ensembles of predictions using Monte Carlo dropout. Using the statistics of these we select batches with high information content for labeling. We show that DEWDROP (1) improves model training efficiency through batch optimization outperforming baselines, and (2) selects data with relevant high information content.

## 1. Introduction

Computational drug design is a key part of any drug development effort [20]. Development of accurate and scalable models for drug design is significantly hindered by the need for large amounts of labeled training data. This challenge arises from the complexity of the models and the vast moleculr design space they must cover. The need to collect such data is costly, time consuming and requires extensive experimental resources[6]. Several methods have been developed and employed to reduce the need for labeled data in computational drug design. Transfer learning[25], data augmentation[5], and foundational models ([9], [17], [1]) are just some of the approaches that have been explored. However, each is limited to specific cases and so cannot fully address the need. Transfer learning relies on the use of drugs for similar targets to the one being studied which are not always available for first in class drugs. Data augmentation relies on very strong assumptions which usually do not hold. Foundational models are powerful but still require labeled data for fine tuning. Active Learning (AL)[8] has also been proposed as a possible way to reduce required data. AL is more general than the other methods and while it relies on an underlying, task specific, model, several AL method can be generally applied in sequential or batch setting[22].

Traditional computational drug design workflows operate in iterative cycles: molecules are tested, results are used to revise the model and a new set of molecules, which according to the model will lead to the best performance, are selected and tested. This process continues until the model achieves satisfactory performance. AL differs from the standard approach in the way it selects the molecules to test in each round (see Figure 1). AL selects molecules not based on their likely good performance but rather based on their ability to improve the model[8,7]. This step often requires the estimation of model uncertainty. The more uncertain the model is about a specific input, the more information we gain from knowing the label for this sample.

**Fig. 1:**
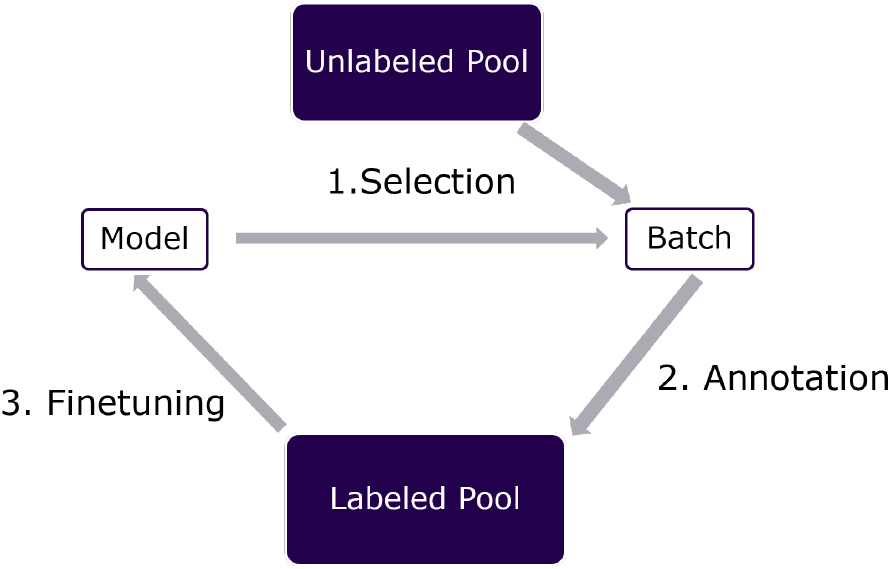
The active learning loop applied to candidate selection.

Several methods have been proposed for using AL in biomedical research. [26] introduces a deep active learning framework that models interatomic potentials by using gradient features from a neural network’s neural tangent kernel to measure data similarity for data selection. [10] applies several active learning strategies to chemogenomics and proteins to learn compact models for protein and ligand interactions. [23] and [13] investigates the effectiveness of active learning selection on both machine learning and deep learning models to predict relative binding free energy (RBFE). While these previous approaches have shown promising results in classification and regression tasks with single or few target variables (for example interaction and free energy calculations), AL for high- or variable-dimensional regression, such as protein structure prediction, has not been successfully achieved with these methods.

Diversity is another important property of good AL selections. This is often required to avoid local minima, especially for batch AL [14]. For example, [3] introduces Batch Active learning by Diverse Gradient Embeddings (BADGE), an active learning strategy that use an estimate of Fisher information as surrogate for model uncertainty, clusters gradients from the last layer of a deep neural network and select the cluster centers as the new batch. Additionally, [2] improves upon BADGE by proposing Batch Active learning via Information Matrices (BAIT), which measures the complete Fisher information matrix by considering the gradient embedding of all samples in the pool. However, BAIT is defined only for fixed-dimensional outputs, and so cannot be used for the data we discuss here. BADGE has demonstrated strong performance on image datasets, but its effectiveness on protein structures has not been evaluated. In [4], a novel active learning method, COVDROP, is shown to outperform baselines in 1D-regression tasks in drug design. Similar to BAIT, such approach is not appropriate for labeling of protein structures.

To address the aforementioned challenges and enable the use of AL in protein structure prediction, we developed DEWDROP and integrated it with Equifold[16], an *SE*(3)-equivariant protein structure model, which was fine-tuned on VHH structures. Unlike BADGE and BAIT, DEWDROP estimates information content by calculating the joint entropy across all samples in the pool using ensembles of prediction generated by Monte Carlo dropout[12]. A batch of samples is then selected through an iterative batch optimization algorithm, which maximizes the joint entropy.

We evaluate the effectiveness of DEWDROP by conducting retrospective experiments on two different classes of proteins: VHH-only and mycobacterial proteins, simulating an active learning environment. The results show that DEWDROP outperforms baseline strategies (Random, K-Means[18], and BADGE) in labeling efficiency. Additionally, we performed candidate selection on VHH sequences and demonstrated that DEWDROP selects batches that maximize uncertainty while maintaining diversity compare to other strategies.

We chose to work with a modified Equifold because of its relative simplicity. It was competitive with SOTA at publication at a fraction of the size and cost, without MSA or any features other than the sequence, and with a relatively straightforward architecture. As noted above, our main focus is on molecules and structures from underexplored classes. For such datasets we can fine-tune a complex model though to do that we need initial labeled data. For this we can use a proxy model like Equifold to select initial sets from the class. Alternatively, DEWDROP can be directly combined with more complex models, at a higher cost in compute.

## 2. Method

We propose a new ensemble-based selection strategy, DEWDROP. The DEWDROP strategy (see Figure 2) selects a new batch of data by maximizing the estimated information content of that batch. For 3D structures, we can estimate joint entropy between their coordinate predictions, as we discuss in Sections 2.3 and 2.3. However, given the potentially large numbers of proteins in each batch, and the large number of different candidate batches considered, it is computationally prohibitive to precisely compute the joint entropy of whole batches of proteins and their coordinates together as a system, each time we need them, even assuming a normal distribution. Therefore, we use an approximation to estimate the entropy of a batch of candidates from the more-precisely-calculated pairwise joint entropies.

**Fig. 2:**
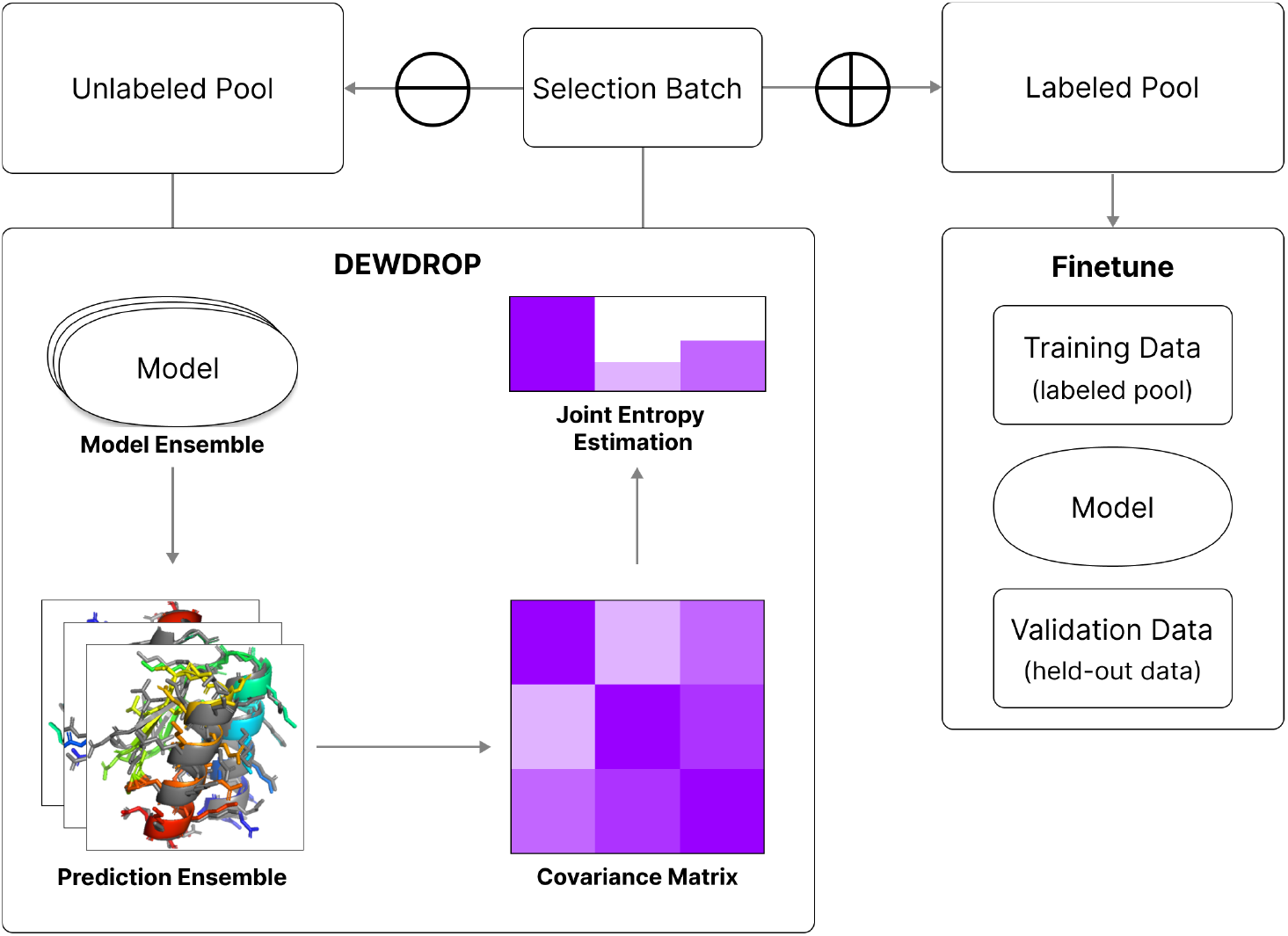
Illustration of the DEWDROP workflow. DEWDROP utilize model ensemble to produce an ensemble of predictions, which is used to produce the covariance matrix and subsequently the joint entropy matrix. When DEWDROP select a batch, that batch is removed from the unlabeled sample pool and added to the labeled sample pool after being sent to expert for labeling. Then we use the labeled data, at any selection rounds, for model fine-tuning.

### 2.1 Higher-order independence assumption and approximating the entropy

First we will describe the approximation we will use for the entropy of a batch of candidates. Recall that the entropy of a probability distribution *P* over a measure space *X* is

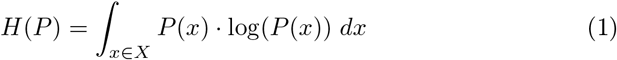

If we have *N* random variables, their joint entropy is just the entropy of their joint probability distribution:

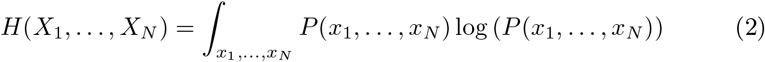

The *dual total correlation* of *n* random variables *X*_1_, …, *X*_*n*_ is

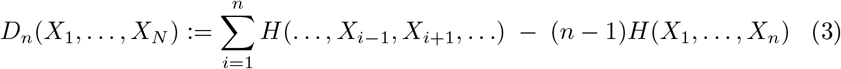

Therefore, for a set of *N* random variables, we say the dual total correlation vanishes at order *k* if *D*_*k*_ (𝒳) = 0 for all order-*k* subsets 𝒳 ⊂ {*X*_1_, …, *X*_*N*_,}, |𝒳| = *k*. If the dual total correlation vanishes at all orders for a set of variables, those variables are *independent*. Rather, we will assume that dual total correlations greater than 2 are *small*. This allows arbitrary correlations between candidates, but no *higher-order correlations*.

#### Assumption

We assume that dual total correlations vanish for *k* > 2 (or approximately so). In this case, an argument by induction gives entropy in terms of pairwise joint entropies:

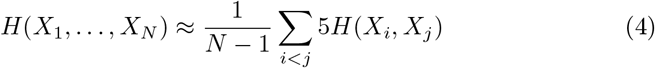

Therefore, we use the ensemble of predictions to estimate pairwise joint entropies *H*(*X*_*i*_, *X*_*j*_), and we seek batches that maximize the acquisition function:

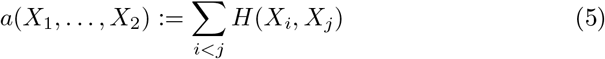

### 2.2 DEWDROP for discrete targets

As mentioned above DEWDROP generates prediction ensembles, estimates the information content with a joint entropy matrix, and selects new batches for labeling through a batch selection process that optimizes the objective described in Equation (5).

We first discuss computation of joint entropy’s for the discrete case. To obtain the ensemble of predictions for each selection round, we inject *M* dropout seeds to simulate different sets of parameters, where *M* is the size of the ensemble. This is a computationally cheap yet effective approximation of model uncertainty [12]. Formally, the predictions form an ensemble, **E** ∈ ℝ^*N* × *M*^, over all samples from the unlabeled pool, 𝒰 = {*x*_0_, …, *x*_*N*_}, where *N* is the size of the unlabeled pool and *x*_*i*_ ∈ ℝ^# residue×3^. An adequately large value for *M* is crucial for a good approximation; however, the choice of *M* is upper-bounded by the computational resources available.

We define target labels as *K* = {0, …, *k* ™ 1} and an ensemble predicted target 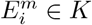 for every prediction in an ensemble set **E**. For any two predicted targets, 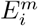 and 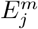, we calculate the joint entropy matrix, *H*_*ij*_, as:

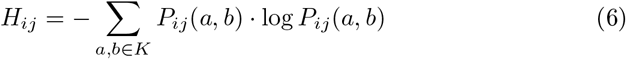

where *P*_*ij*_ is the joint probability distribution that 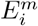 is label *a* and 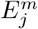 is label *b*, which is formally defined as follows:

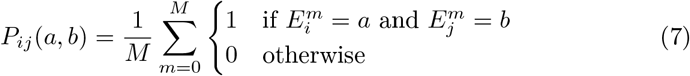

DEWDROP employs an iterative optimization algorithm to select the batch with the highest net joint entropy. The algorithm begins by defining a probability distribution over all samples, based on their self-entropy values. It then starts with a randomly initialized batch and computes the acquisition function as described in Equation 5. Over a set number of iterations, the batch is refined by systematically replacing one sample with unselected candidates from the pool. For each new configuration, the acquisition function is recalculated, and the candidate that maximizes the score improvement is selected. The pseudocode for this batch optimization algorithm is provided in the supplementary material.

### 2.3 DEWDROP for continuous targets

We next extended the DEWDROP framework to the problem of predicting atom coordinates, which is a regression problem. We made the following assumption to accommodate the continuous variables as targets: the probability density function of the coordinates of each group representation of atoms from candidate *i* follows a normal distribution, i.e. 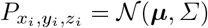. Therefore, the distribution is completely characterized by its mean and covariance (which we compute empirically, see Section 2.3). Using our assumption, we can redefine the joint probability density function of two predictions for two candidates, *i* and *j*, as another multivariate normal distribution:

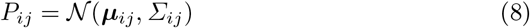

where ***µ***_*ij*_ is the multivariate mean, and *Σ*_*ij*_ is the covariance matrix, both of which are calculated with respect to the predicted coordinates of every group representation in both candidates *i* and *j*.

We then calculate the joint entropy matrix of any two predictions, denoted as 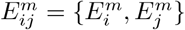, following the joint entropy formula for continuous random variables:

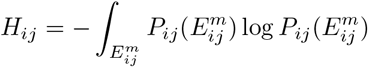

We start by expanding log 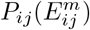 into terms composed of parameters of the multivariate normal distribution ***µ***_*ij*_, *Σ*_*ij*_, and the input 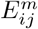:

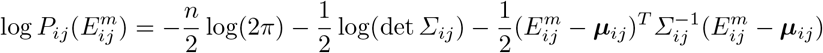

where *n* is the dimensionality of the pair of coordinate predictions 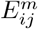. Substituting the new log *P*_*ij*_(*X*_*ij*_) into the joint entropy calculation:

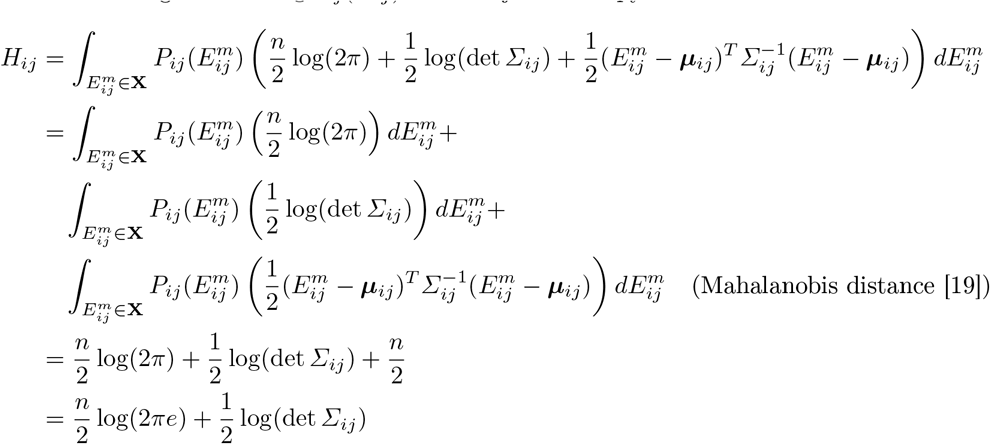

The constant term, 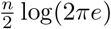, acts as a scaling factor for the determinant of the covariance matrix, det(*Σ*_*ij*_). Since the dimensionality parameter, *n*, remains constant across sample pairs, this term becomes a constant scaling factor, allowing us to ignore it when comparing joint entropy between samples from the same dataset. In addition to the smoothing effect introduced by our high-order independence assumption, we add a small adjustment factor, *ϵ*_**I**_, to the diagonal of the covariance matrix in order to address numerical instability during matrix decomposition. This is equivalent to adding Gaussian noise of variance *ϵ*_**I**_. The final joint entropy calculation is as follows:

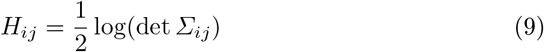

#### Empirical estimation of covariance

For our experiments, we estimated the covariance of our predictions in the following way. We trained with dropout layers, using them to approximate the posterior distribution as in [12].

We generated a correlated ensemble of 128 predictions for each position in each protein, we aligned the predicted structures using the Kabsch-Umeyama algorithm [24], and reduced this to 128 principal components. We then used this ensemble of predictions to estimate the total covariance between any two protein structures, and computed joint entropy estimates as above.

Our choice of ensemble size of 128 was justified by a robustness analysis using bootstrapping to estimate the variance of our joint entropy calculations (see Supporting Methods). 128 samples proved to be more than enough to get consistent rankings in realistic tests.

### 2.4 Datasets

#### Nanobodies

For evaluation and fine-tuning, we collected 1,708 VHH structures from .cif records on the RCSB Protein Data Bank (RCSB PDB) and compiled them into the PDB1700 dataset. All chain sequences were selected from complexes consisting of VH (Variable region of the Heavy chain), VL (Variable region of the Light chain), and Ag (Antigen chain). Each complex contains at least one of these chain types. The chains were validated through self-alignment by comparing the sequence from the file header to the residues from atom coordinate records. Missing residues, as determined by alignment, were marked as UNKNOWN, with NaN used for their coordinates. After numbering with ANARCI [11], the chains were categorized as VH, VL, or Ag and reassembled for comparison to the original complex structures. The PDB1700 dataset includes only VHH structures from VH-Ag complexes.

#### Mycobacteria

For a second benchmarking dataset, we chose another non-human protein class: those found in *mycobacteria*. We used the dataset of the Alphafold-predicted structures for mycobacterial proteins [15]. We filtered to a maximum size of 500 residues, giving us 1335 proteins in total. Limiting our pool to smaller proteins allows a smaller Equifold models with fewer parameters to perform well on limited compute resources, while also enabling a performance comparison between DEWDROP and other baselines. This means that our second dataset is half-synthetic (real sequences, predicted structures). We chose it because it covers a non-animal, non-plant species (which are generally not well-covered by experimental structures), and has the desired size of order 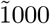.

## 3 Results

We evaluated the DEWDROP active learning framework on on two datasets— PDB1700 VHH (nanobodies) and a Mycobacteria dataset—by conducting a retrospective experiment, simulating a fixed number of AL iteration rounds, and comparing the validation performance of the protein structure model at each round. In addtion, we assessed the quality of the selected batches for each strategy by analyzing the model’s perceived uncertainty, quantified by joint entropy, and the actual uncertainty, measured by validation loss, associated with each selected sample.

### 3.1 Retrospective Experiments

#### Nanobodies

We first evaluated Dewdrop and compared it with prior methods for AL by testing them on a VHH structure prediction task. For this we fine-tuned the Equifold model [16]. The Equifold model is initially trained an antibody dataset from the Structural Antibody Database (SAbDab) [21]. We used PDB1700 VHH dataset to fine tune the model for this molecule type. The resulting model is referred to as Nanofold. The retrospective experiments simulate a real production setting, where dataset labeling and model training occur concurrently, and the improvement in quantitative metrics from training on each batch corresponds to the quality of the batch selected by the active learning strategy.

#### Mycobacteria

In addition, we ran a further retrospective experiment on a mycobacteria dataset. This dataset was composed of 1335 protein structures, and we held back 10% for test and 10% for validation. We used the same setup as with Nanobodies, with exceptions noted below. Our initial weights before active learning was applied were Equifold trained on human antibodies.

#### Comparisons

We compared the DEWDROP selection strategy with several prior methods and a random selection baseline. For the prior methods we tested those that can be readily applied to large regression problems (BADGE and K-Means [18]). Our simulations proceeded in batches for all methods. For each batch, we used these methods to select 100 molecules without referencing their ground truth coordinates. Separate versions of the model were trained with the batches selected by different strategies, and the final validation metrics were recorded. The pseudocode for the retrospective experiment is shown in the supplementary material.

We ran the experiment similarly but not identically for the two datasets:

- For the PDB1700 VHH dataset, we conducted 6 batch selection rounds for each trial, and fine-tuning for each round was terminated when (1) the model had trained for 200 epochs, or (2) the model converged. We ran 3 trials for each method.
- For the mycobacteria dataset, we reduced the number of selection rounds to 4, since later rounds did not prove to be more informative with the first dataset. We ran 3 trials for each of DEWDROP, K-means and Random selection.

#### PDB1700 VHH results

Figure 3 shows the average of three loss metrics across all trials for *PDB1700 VHH* : frame-aligned point error (FAPE), angle loss, bond loss, and the aggregate of all three losses called total loss. From Table 8a and 8b, we see that DEWDROP after the first round of selection have shown a clear advantage over other strategies in FAPE loss, and a better total loss as a consequence. Both FAPE and total validation loss indicates that DEWDROP-selected batches had higher information content, as models trained on these batches reached the target loss threshold faster than the baseline. However, for the angle and bond loss (see table 8c and 8d), BADGE is doing the best with DEWDROP performs second to BADGE. One explanation is DEWDROP only considers the predicted 3D coordinates of the atom representation, so it is selecting candidates which have the most uncertainty in positional alignment instead of angle or bond length. BADGE, on the other hand, looks at the last layer gradient that updates all parameters which including the partial gradient with respect to angle and bond loss, thus there should be more information about angle and bond configuration using BADGE than other strategies.

**Fig. 3:**
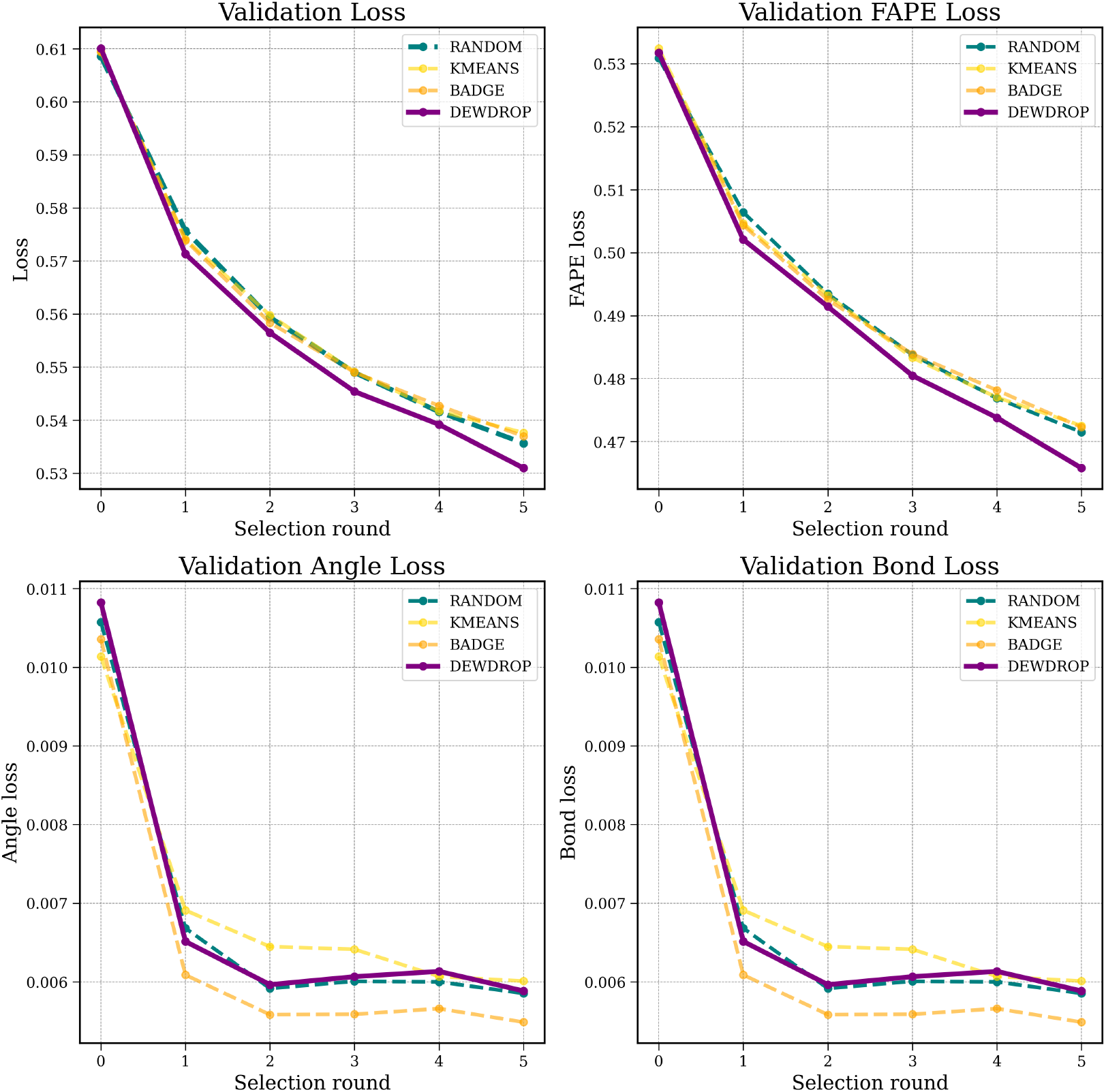
*Nanobodies* – Validation metric across different selection rounds for DEWDROP, BADGE [3], K-Means [18], and Random strategy for the PDB1700 VHH dataset. The loss values are averaged across all trials. From the top two graphs, we see DEWDROP achieve lower structural loss value faster than other baselines. However, BADGE is superior than DEWDROP and others in the angle and bond loss performance.

#### Mycobacteria results

Figure 4 shows loss metrics across all trials for the *mycobacteria* dataset: total validation loss, angle loss, bond loss, and frame-aligned point error (FAPE). From Table **??**, we see that DEWDROP outperforms Random and Kmeans baselines, with a clearer advantage after the second selection round. In the first round, where limited information is available, the performance gap is smaller, but by the second round, DEWDROP-selected batches lead to models that reach the target loss threshold faster. This trend is consistent across the different loss metrics (with the exception of angle loss, as with the first dataset), with differences beyond the standard error, indicating that DEWDROP-selected batches contain higher information content. Note that, in this case, FAPE loss hardly changes in absolute terms from start to finish. A likely explanation is that the pretrained, starting model was already well-adapted to minimize FAPE on this dataset.

**Fig. 4:**
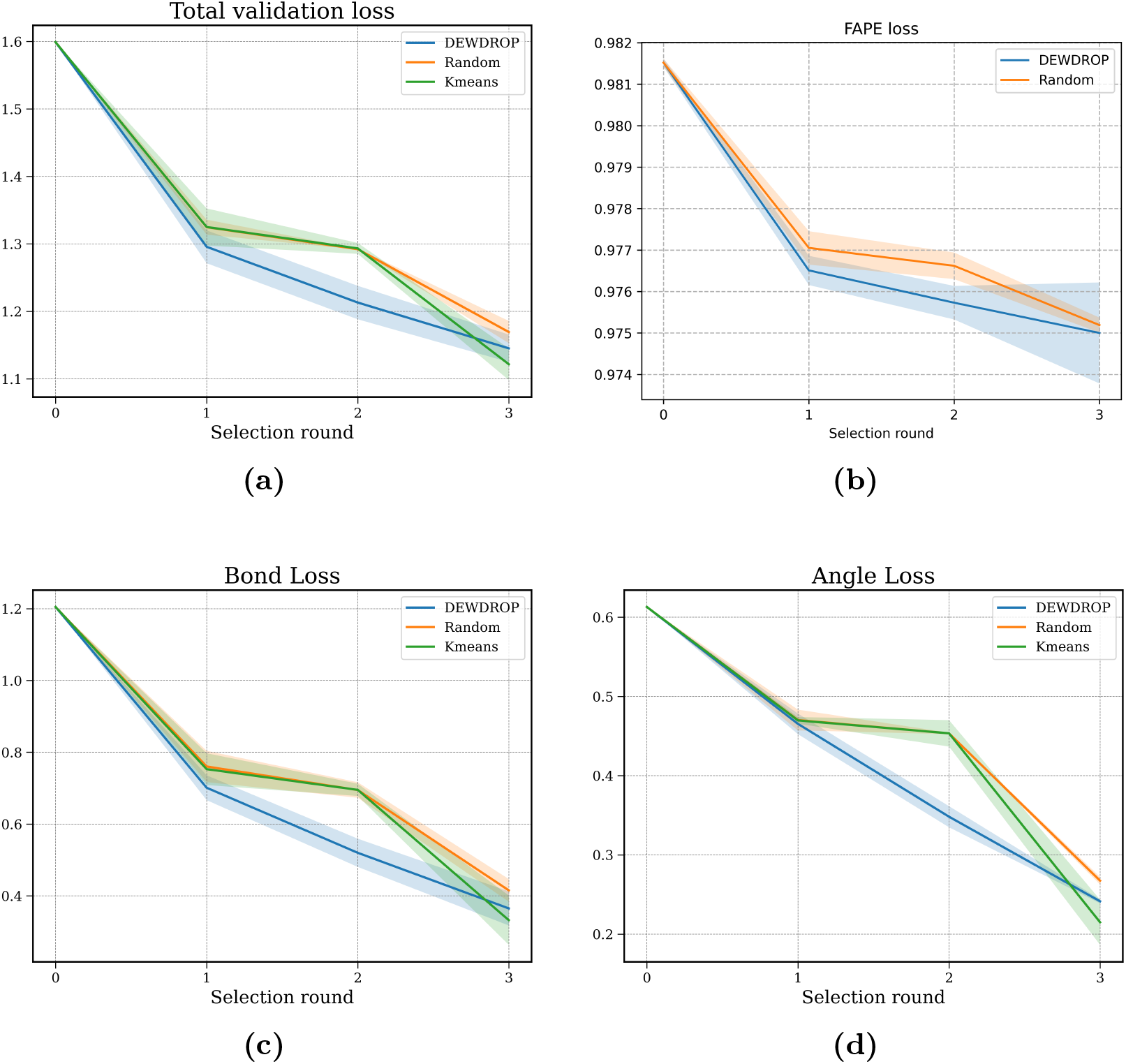
*Mycobacteria* Validation metric across different selection rounds for DEWDROP, K-Means [18], and Random strategy for the Mycobacteria dataset. The loss values are averaged across all trials, with standard errors shown. We see DEWDROP achieve lower loss value faster than other baselines.

### 3.2 Batch Selection

To assess DEWDROP’s ability to select highly uncertain samples that could maximally enhance model performance, we compared the quality of the batch selected by DEWDROP with the batch selected with BADGE and K-Means. Specifically, we chose 100 VHH sequences using K-Means (diversity-based), BADGE, and DEWDROP selection strategies. Then, we evaluated the combined structural loss of each predicted VHH structure compared to the ground truth, alongside the model’s uncertainty on those selections, which was quantified using their joint entropy relative to the rest of the dataset. The results are visualized in Figure 5.

**Fig. 5:**
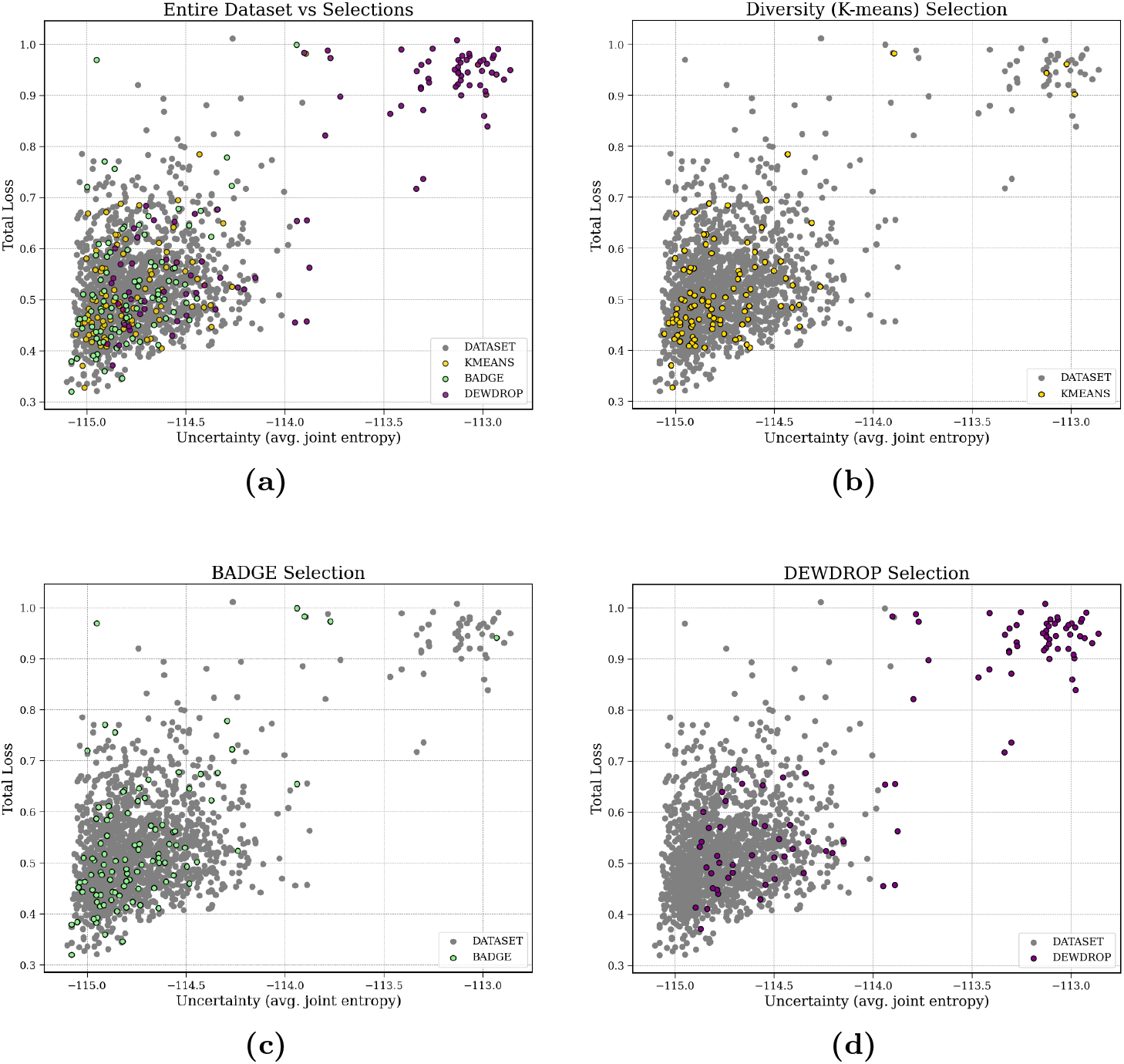
Comparison across different selection strategies: total validation loss of predicted structure over model uncertainty (average joint entropy). 5a compares all 4 strategies with the entire dataset. 5b, 5c, and 5d compares each strategy’s selection (colored) with the distribution of the entire dataset (grey). The horizontal axis represents uncertainty, with higher uncertainty on the right and lower uncertainty on the left.

**Fig. 6:**
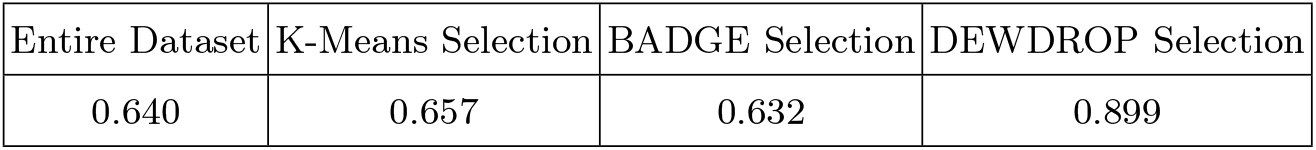
Pearson correlation coefficient between combined structural loss and average joint entropy. K-Means and BADGE strategy select batches with similar correlation as the dataset in their measured uncertainty and total validation loss. DEWDROP select a batch with significantly higher correlation compare to the dataset.

The results indicate a clear distinction between the batches selected by DEWDROP, BADGE, and K-Means. K-Means selects samples that reflect the overall data distribution, which can ignore valuable samples with high uncertainty. BADGE also includes very few samples with high uncertainty. In contrast, the DEWDROP selected batches a good balance between representative and uncertain samples. Roughly half the selected set is coming from the densely populated (but well resolved) samples whereas the other half is focused on the most uncertain samples. This leads to faster convergence while still being able to capture dense regions in the input space. Furthermore, we computed the Pearson correlation between structural loss and average joint entropy, revealing a strong correlation between estimated model uncertainty and the observed error in the DEWDROP-selected batch (Table 6). This correlation further establish joint entropy as an accurate estimation of the true model uncertainty, proving the DEWDROP strategy is capturing an effective subset for model training.

## 4 Discussion

DEWDROP outperforms the diversity-based criteria in total loss and FAPE loss, but is in turn outperformed in angle prediction. We believe the reason for the relative underperformance of DEWDROP in angle loss is that we use a global alignment on the ensembles of predictions; i.e., all 128 predicted structures for a given protein are aligned with each other globally, using only rigid transformations. While this is a good measure of large-scale variance, it’s quite possible for a protein structure prediction to be accurate in the large scale while having small-scale irregularities which produce large angle errors.

We can see in Figure 5 that DEWDROP’s selection differs from the other selections in that it favors uncertain candidates for selection. However, it doesn’t only select the N most uncertain. It has a kind of diversity criterion too. If two candidates have uncertainty which is highly correlated, then their mutual information will be high, and their paired joint entropy will be correspondingly less than the sum of their entropies. If correlation is taken to be a measure of similarity, then this will tend to favor batches of dissimilar candidates.

We can see that this interpretation of correlation makes some sense in Figure 7. We see, using the same last-layer embedding space as with K-means and BADGE, that lower distance in the embedding space tends to yield higher correlation scores. So these two different measures of similarity/diversity approximate each other.

**Fig. 7:**
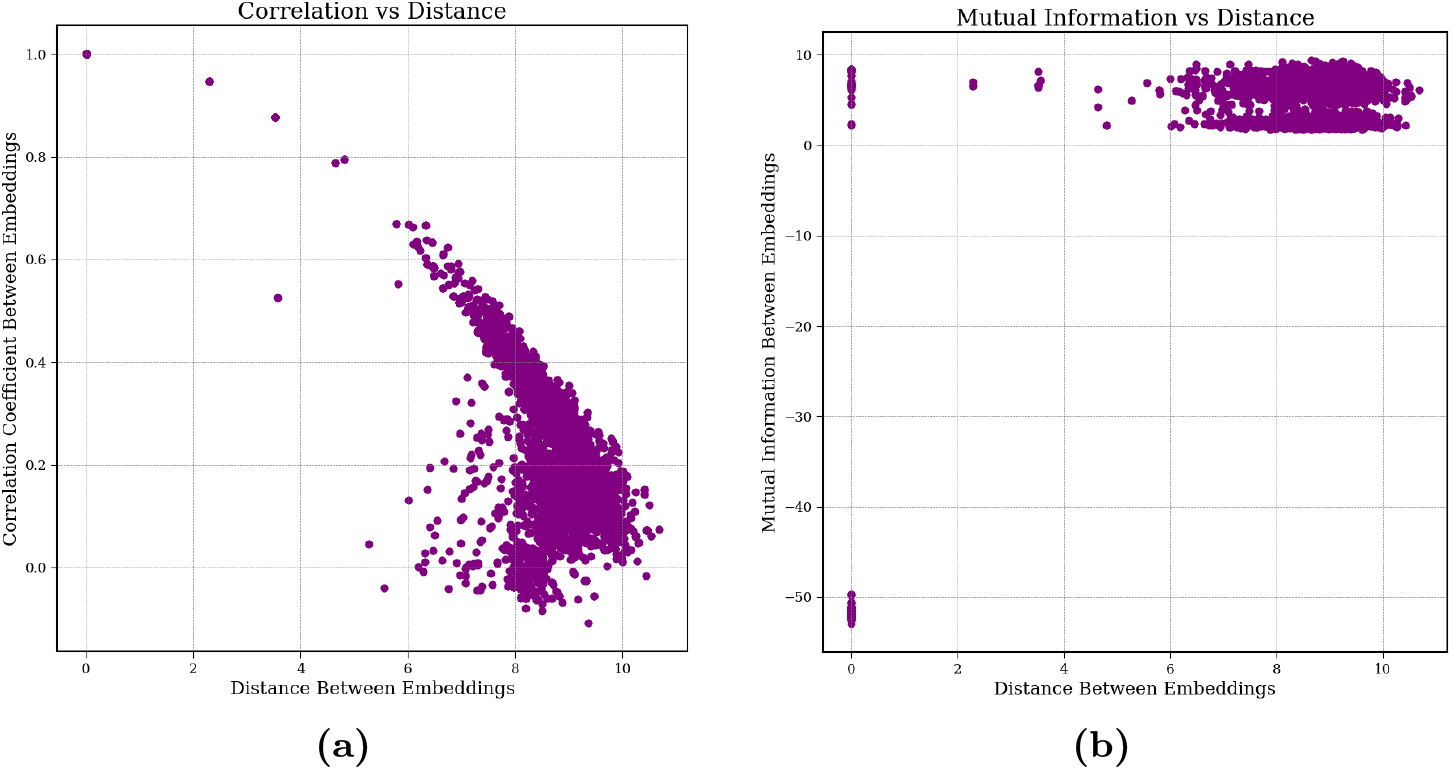
We investigated the properties of the embeddings extracted from the last layer for each output structures by examining their distance, correlation coefficient, and mutual information. (a) shows the correlation between the correlation efficient and the distance between embeddings. (b) compares the same distance between embedding against their mutual information.

**Fig. 8:**
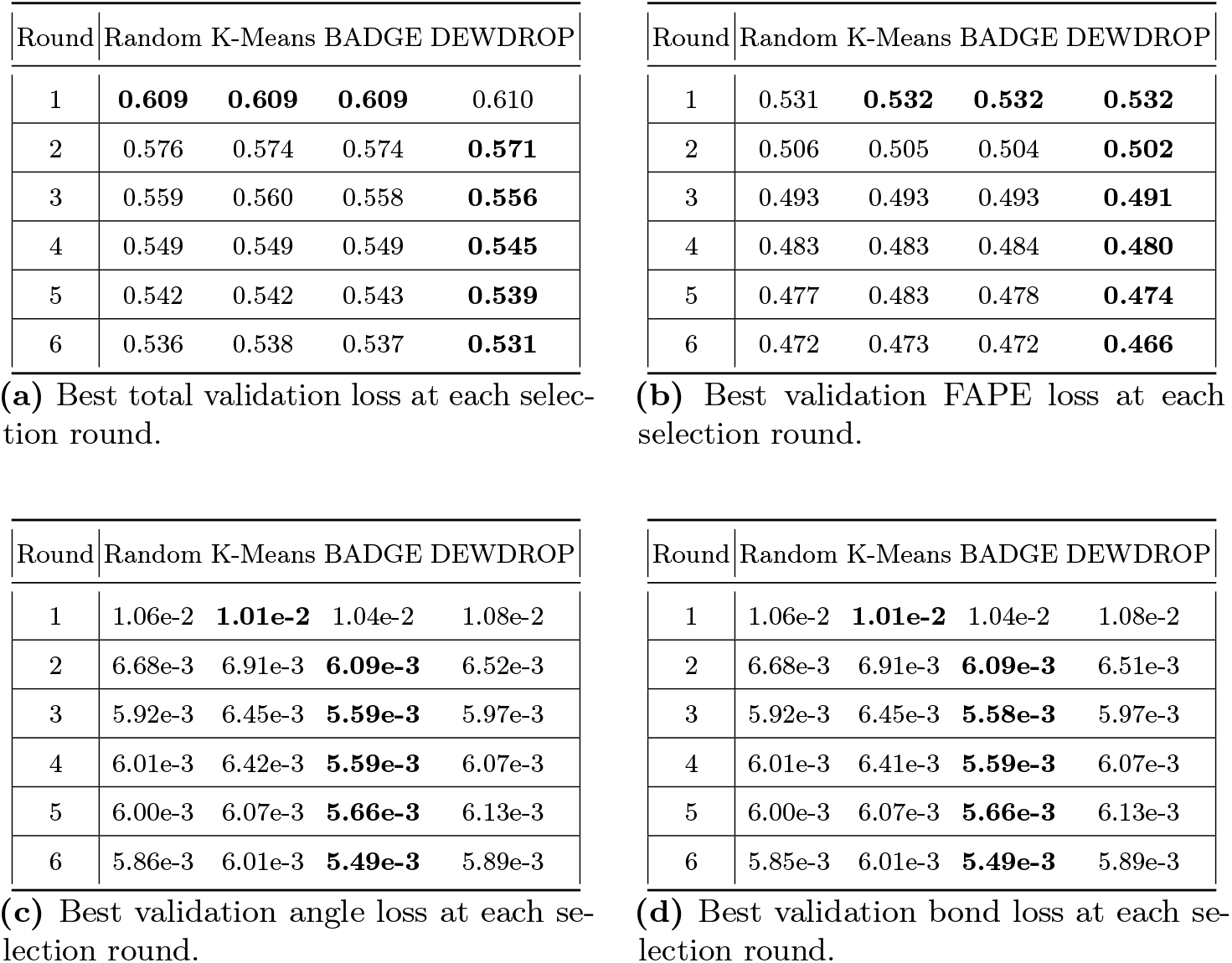
Different types of loss values associated with the validation loss graphs.

This can fail, and the selection can be less diverse than would be ideal, if correlations are being systematically underestimated. While we can see from Figures 5 and Table 6 that the estimates of the uncertainties of individual candidates are fairly good. But estimating correlations is a tricky problem, and any weaknesses in the method will tend to add uncorrelated noise.

If the estimated correlations were higher, we would expect that DEWDROP would select more sparsely from the upper-right cluster of highly uncertain candidates, and some of those slots would instead go to other candidates, presumably giving better diversity coverage, and removing one of BADGE/K-means’s advantages. Indeed, one could modify DEWDROP to artificially increase correlations, and this seems a promising avenue for further development.

## 5 Conclusion

In this work, we propose a novel active learning strategy, DEWDROP, for protein structure prediction. DEWDROP, unlike BADGE, measures model uncertainty through a joint entropy calculation, directly using an ensemble of model predictions. Our results demonstrate that DEWDROP outperforms baseline strategies, including Random, BADGE, and K-Means, by achieving superior evaluation metrics in fewer selection rounds. A comparative analysis of batch selection demonstrates that DEWDROP prioritizes sequences with high model uncertainty, rather than merely reflecting the overall data distribution, unlike K-Means and BADGE.

These findings establish DEWDROP as a state-of-the-art active learning strategy for protein structure prediction, while remaining flexible with the choice of model. This presents a key advantage over other approaches that rely on gradients or embeddings, which are challenging to extract from complex deep neural networks like Equifold. Additionally, the strong correlation observed between structural loss and uncertainty estimation highlights a promising avenue for developing new uncertainty heuristics in future research.

## 6 Supplementary Material

### 6.1 Pseudocodes

#### 1 Algorithm

Batch optimization

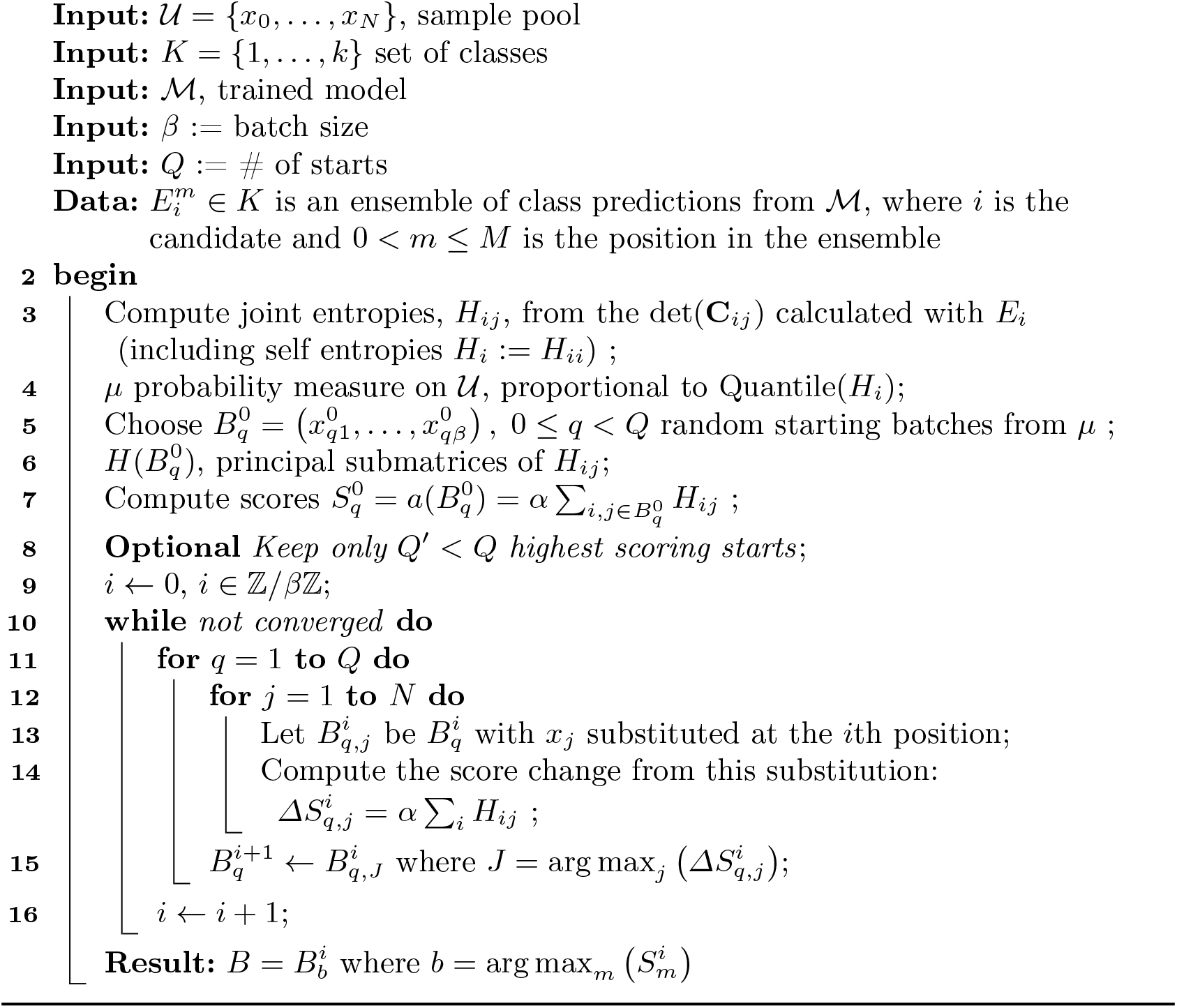

#### 1 Algorithm

Retrospective experiment

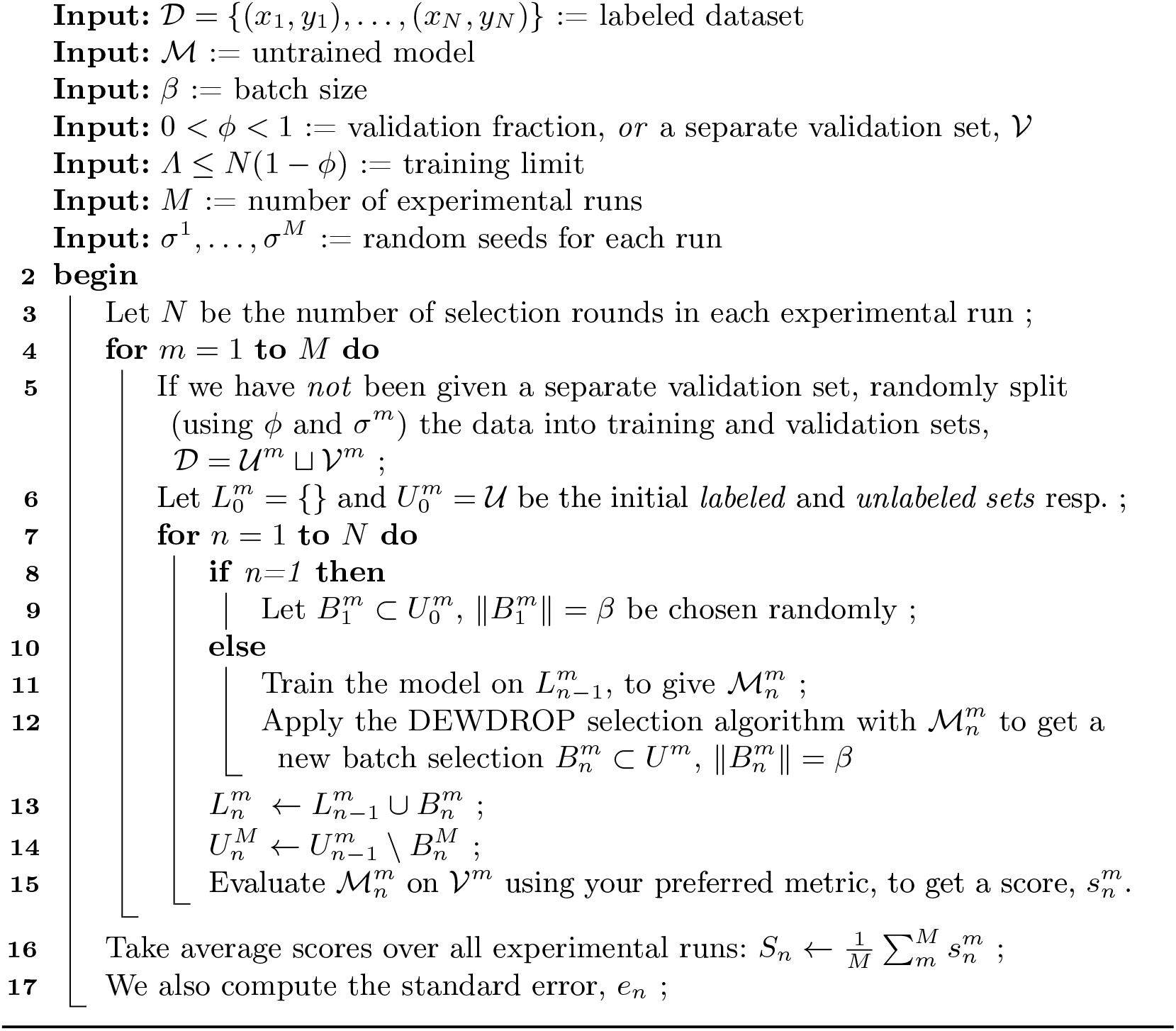

### 6.2 Detail Results for the Retrospective Experiments

Project GitHub: https://github.com/Sanofi-Public/dewdrop

## Notes

### Competing Interest Statement

The authors have declared no competing interest.

